# Lesions to different regions of frontal cortex have dissociable effects on voluntary persistence

**DOI:** 10.1101/2023.11.16.567406

**Authors:** Camilla van Geen, Yixin Chen, Rebecca Kazinka, Avinash R. Vaidya, Joseph W. Kable, Joseph T. McGuire

## Abstract

Deciding how long to keep waiting for uncertain future rewards is a complex problem. Previous research has shown that choosing to stop waiting results from an evaluative process that weighs the subjective value of the awaited reward against the opportunity cost of waiting. In functional neuroimaging data, activity in ventromedial prefrontal cortex (vmPFC) tracks the dynamics of this evaluation, while activation in the dorsomedial prefrontal cortex (dmPFC) and anterior insula (AI) ramps up before a decision to quit is made. Here, we provide causal evidence of the necessity of these brain regions for successful performance in a willingness-to-wait task.

28 participants with frontal lobe lesions were tested on their ability to adaptively calibrate how long they waited for monetary rewards. We grouped the participants based on the location of their lesions, which were primarily in ventromedial, dorsomedial, or lateral parts of their prefrontal cortex (vmPFC, dmPFC, and lPFC, respectively), or in the anterior insula. We compared the performance of each subset of lesion participants to behavior in a control group without lesions (n=18). Finally, we fit a newly developed computational model to the data to glean a more mechanistic understanding of how lesions affect the cognitive processes underlying choice.

We found that participants with lesions to the vmPFC waited less overall, while participants with lesions to the dmPFC and anterior insula were specifically impaired at calibrating their level of persistence to the environment. These behavioral effects were accounted for by systematic differences in parameter estimates from a computational model of task performance: while the vmPFC group showed reduced initial willingness to wait, lesions to the dmPFC/anterior insula were associated with slower learning from negative feedback.

These findings corroborate the notion that failures of persistence can be driven by sophisticated cost-benefit analyses rather than lapses in self-control. They also support the functional specialization of different parts of the prefrontal cortex in service of voluntary persistence.

## Introduction

In order to achieve desirable outcomes, we must often devote time towards a goal. From mastering a new skill to developing meaningful relationships, many realms of human activity benefit from perseverance. This notion has led to the conventional wisdom that grit is a key to success and personal development.^1^ In a laboratory setting, delay of gratification paradigms like the well-known marshmallow test^2–4^ have measured persistence by evaluating how long people— mostly children—are willing to wait to receive a desirable reward. Individual differences in the ability to delay gratification in these tasks correlate with lifelong success, and longer wait durations are taken to be indicative of greater discipline and self-control.^5–7^

That being said, there are also situations in which persistence leads to suboptimal outcomes. For example, a test taker who gets stuck on a question and devotes the rest of the time to solving it will most likely get a bad grade overall. Similarly, a tourist who waits in line for hours for the city’s most popular chocolate chip cookie is probably missing out on more exciting sight-seeing experiences. Thus, although grit and perseverance can be necessary for success, they can also be misapplied – any attempt to evaluate the optimality of committing time toward a prospective reward must take the broader decision context into account.^8–10^

In fact, many psychiatric disorders are characterized by either too much or too little persistence. For instance, obsessive-compulsive disorder (OCD) often involves repeated performance of rituals that are time-consuming and not causally related to tangible success.^11,12^ Conversely, depression can lead people to give up sooner than they should, even when the outcome is valuable and approaching quickly.^13^ Individuals who struggle with addiction often have difficulty sticking with the choice of the delayed but ultimately more fulfilling rewards associated with sobriety, especially when faced with the immediacy of the drug. A more comprehensive understanding of how people assess when persistence is worthwhile and when it is not will pave the way for clinical interventions that may help improve decisions.

Previous research has shown that healthy adults do indeed leverage the contextual features of their environment in order to calibrate persistence.^14–19^ This adaptability has most explicitly been demonstrated in the realm of waiting behavior, when participants must decide how long they are willing to wait for an uncertain future reward. Thus, although persistence can take many different forms, including active effort expenditure, we will focus here on willingness to wait.

We have previously shown that people use the temporal statistics of reward delivery to adaptively determine whether to persist.^14,15^ In those experiments, participants waited for a monetary reward in two temporal environments: one in which the distribution of wait durations was heavy-tailed and another in which it was uniform. In the heavy-tailed condition, the expected time remaining until the reward arrived increased the longer one had already waited. The reward-maximizing behavior was therefore to move on to a new trial after a short delay. Conversely, in the uniform condition, the expected time remaining until the reward arrived decreased as one waited, meaning that the optimal behavior was to continue waiting as long as necessary. Participants’ behavior paralleled these normative principles, as they displayed significantly greater willingness to wait in the uniform environment. In general, performance on these sorts of willingness-to-wait (WTW) tasks reflects the results of a dynamic and context-dependent comparison between the anticipated value of the potential reward and the opportunity cost of waiting for it.

Neural evidence further corroborates this framework: McGuire & Kable^15^ found that activity in the vmPFC, an area of the brain sensitive to subjective value^20–22^, evolved differently in the different temporal environments. If the distribution of wait durations favored persistence (uniform distribution), the vmPFC signal ramped up progressively as time passed and the wait shortened. Conversely, if the distribution of wait times was such that the value of waiting decreased after a certain point (heavy-tailed distribution), activity in the vmPFC was flat until people chose to quit (see ^23^ for a parallel finding). Furthermore, activity in several brain regions – including the insula and dorsal regions of the prefrontal cortex –ramped up before the decision to quit was made.^15^ Beyond providing insight into the neural mechanisms that underlie persistence in willingness-to-wait tasks, these findings are important because both the location and time course of neural activity suggests a dynamic assessment of subjective value.

Although these fMRI results provide evidence that the vmPFC, dmPFC, and anterior insula are involved in adaptive persistence, they cannot demonstrate that these brain regions are *necessary* for such behavior. To address this question, we collected behavioral data in a sample of 28 participants with lesions to different areas of the frontal cortex, as well as in 18 control participants without brain lesions. Doing so allowed us to assess whether damage to regions known to be engaged during this task cause deficits in people’s ability to calibrate persistence. We separated our sample into 3 groups based on the location of the lesion: one group (n = 8) had lesions that mostly affected the dorsomedial prefrontal cortex (dmPFC) or anterior insula (AI), another group (n = 10) had lesions to the vmPFC, and the final group (n = 10) had lesions in other areas of the frontal cortex (frontal controls). Behaviorally, we found that participants with lesions to vmPFC waited less in both reward-timing conditions, while participants with lesions to the dmPFC/AI were significantly impaired in calibrating their waiting behavior across conditions. We found no evidence for any impairment in the frontal controls.

Finally, to better understand where in the decision-making process participants with lesions differed from controls, we fit a newly developed computational model of adaptive persistence.^24^ The model tracks running estimates of the value of waiting at each time step within a trial, and probabilistically predicts waiting behavior as a result. It includes 5 participant-specific parameters that describe the learning process – two parameters that capture the speed of learning from positive (α) or negative (ν) feedback, respectively; an inverse temperature parameter (τ) that determines action selection noise; a discount parameter (γ) for temporally remote rewards; and a parameter that captures participants’ initial willingness to wait (*η*). We found that lesion location was related to dissociable deficits in the learning process: whereas participants with lesions to the vmPFC exhibited a lower initial willingness to wait (lower *η*), participants with lesions to the dmPFC/AI learned more slowly from negative feedback (lower ν). Taken together, these findings suggest that regions of frontal cortex make computationally distinct contributions to adaptive persistence.

## Materials and Methods

### Participants

A total of 18 healthy controls and 31 participants with brain lesions were recruited from the University of Pennsylvania Focal Lesion Database and the McGill Cognitive Neuroscience Research Registry to participate in this study. The lesion group and controls were matched in terms of age (mean age = 57 ± 9 years for the controls versus 59 ± 12 years for the participants with lesions) and years of education (mean = 14.3 ± 2.1 for the controls versus 14.5 ± 2.5 for the lesion group). Lesions were drawn on a common neuroanatomical template (Montreal Neurological Institute) by neurologists at the research sites who were not aware of task performance.

The results from McGuire & Kable^15^ suggest that different regions of the frontal cortex underlie specific aspects of willingness to wait behavior. Consequently, we separated the participants into subgroups based on the location of their lesions – vmPFC, dmPFC, AI, or elsewhere in the frontal lobe. To establish these groupings, we first created an anatomical mask of each region using the Automated Anatomical Labeling (AAL) atlas^25^. For the vmPFC mask, we included parcels 23–28, 31 and 32, which we truncated to exclude voxels superior to z = 10. For the dmPFC mask, we included regions 5, 6, 19, 20, 32, 33, and the portions of 23, 24 and 31, 32 that were not part of the vmPFC mask (truncated to exclude voxels at or inferior to z = 10). We created the AI mask by combing parcels 29 and 30 of the AAL.

Having established anatomical masks, we leveraged the findings from McGuire & Kable^15^ to narrow our regions of interest (ROIs) to voxels in which activation was sensitive to different aspects of waiting behavior. We intersected the anatomical masks with masks from relevant functional contrasts. For instance, since McGuire & Kable^15^ found that the anterior insula showed differential activation for “quit” versus “reward” trials, we created our final anterior insula ROI by determining the conjunction between significant voxels from that contrast and the AAL-derived mask. The resulting ROI was used to define the anterior insula group, and any person whose lesion comprised >25% of that ROI was classified as belonging to it. For the remaining 25 participants, we first assessed whether at least 50% of the participant’s lesion affected the frontal lobe, as defined by a combination of regions in the AAL atlas. This resulted in the exclusion of 3 participants whose lesions did not meet this threshold. To establish the vmPFC lesion group, we determined the conjunction between the anatomical vmPFC mask and voxels whose activation was found to be sensitive to the subjective value of waiting in McGuire & Kable^15^. Any of the 22 remaining participants whose lesions overlapped with at least 5% of the resulting vmPFC ROI were then classified as belonging to the vmPFC group. Finally, we identified a dmPFC group, whose lesion overlapped with at least 5% of the intersection between regions of the dmPFC sensitive to “quit” vs. “reward” trials^15^ and the atlas-derived dmPFC mask. Overall, this procedure resulted in 6 people with anterior insula lesions, 10 with vmPFC lesions, and 2 with dmPFC lesions. Because the relevant regions of the anterior insula and dmPFC were both shown to be differentially active for “quit” vs. “reward” trials, we combined the anterior insula and dmPFC lesion groups to result in a combined sample of 8 participants with dmPFC/AI lesions (see Figure 1A). Finally, individuals who met the frontal lobe cutoff but not the cutoff for inclusion in either the vmPFC or dmPFC groups were classified as frontal controls (n = 10).

**Figure 1.**
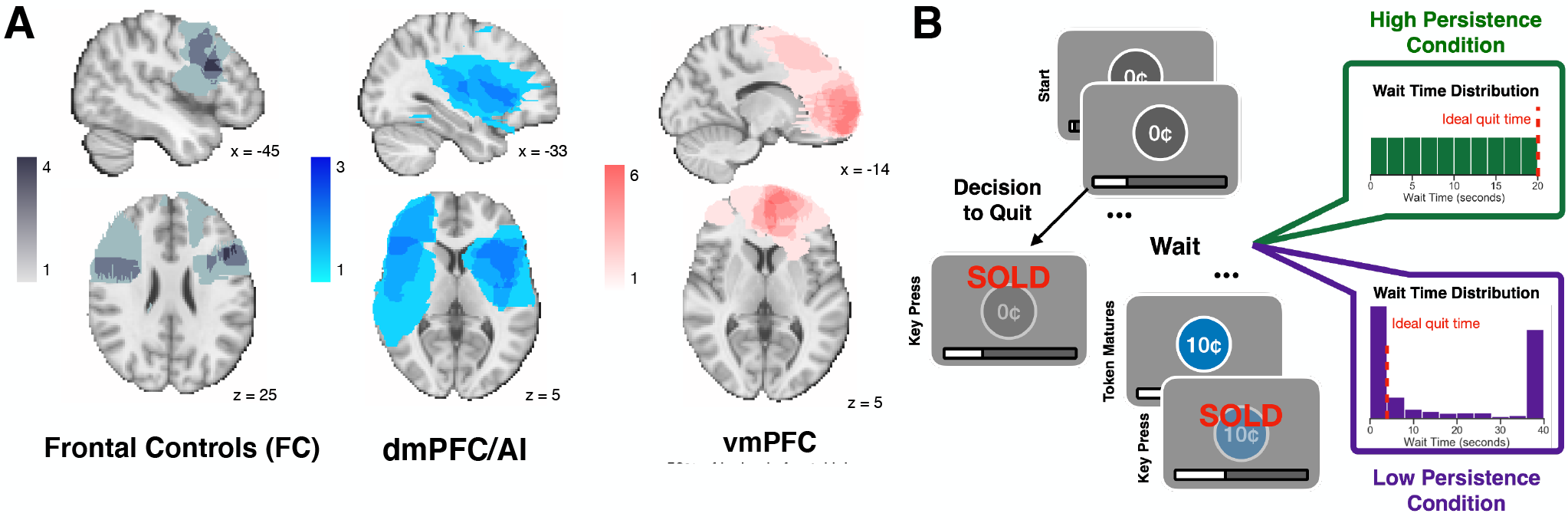
**(A)** Illustration of the degree of overlap in lesion location across participants in each lesion group. **(B)** Overview of the willingness to wait task. Participants chose whether to keep waiting for an uncertain future reward in two temporal environments: one in which the distribution of wait durations was uniform (high persistence condition), and another in which it was heavy-tailed (limited persistence condition).

### Task

In this experiment, we used a willingness-to-wait task similar to the one described in McGuire & Kable^15^ (Figure 1B). Participants were tasked with making as much money as possible in 12 minutes by determining when to sell coins whose value varied within a trial. Specifically, participants received a coin worth 0¢ at the start of every trial. They were told that after a certain amount of time, the length of which varied, the coin would mature and become worth 10¢. At that point, signaled by a change in the coin’s color, they could sell the coin by pressing the space bar, leading to a 10¢ reward. However, participants were also told that at any point in the trial, they could elect to sell the coin, even if it had not yet matured. In that case, they would not receive 10¢, but they would receive the next coin. Whenever participants decided to sell the coin, the word ‘SOLD’ would appear in red in the center of the screen for 1 second. A progress bar on the bottom of the screen indicated how much time remained in the block, out of the total of 12 minutes.

The blocks were defined according to the distribution of wait durations. In the high-persistence block (HP), wait times were sampled according to a uniform distribution ranging from 0 to 20 s. In this condition, the coin was equally likely to mature at any time within the 20-second interval, and the optimal strategy was always to wait until it matured. In the limited-persistence (LP) environment, wait times were sampled according to a heavy-tailed generalized Pareto distribution truncated at 40 seconds. As is illustrated in Figure 1B, the optimal strategy in the limited-persistence environment was to wait only 2.3 seconds for the coin to mature.

### Procedure

The experiment was conducted over two sessions, with the order of the blocks held constant across participants to highlight differences across lesion groups. Session 1 began with a high-persistence block followed by a limited-persistence block. Participants later returned for session 2 (median of 22 days in between sessions), which began with a limited-persistence block followed by a high-persistence block. The first blocks of each session (HP from session 1 and LP from session 2) were taken as instances of “uncontaminated” learning because no task preceded them and are the focus of the upcoming analyses. The second block of each session tested participants’ responses to transitions in the statistical context and will be used in the learning-model fits presented below. The distinction between the two blocks was signaled by a change in the color of the coin, but participants were not explicitly told about the characteristics of the two wait time distributions.

### Behavioral Analyses

To capture waiting behavior, we conducted three main types of analyses in R (version 4.2.3).

First, we used the “coxme” and “survival” packages to estimate survival curves for each lesion group in the two temporal environments. These survival curves estimated the likelihood that a participant would wait until time *t*, assuming the coin had not yet matured. Because trials in which participants waited until reward receipt provided only a lower bound on how long they would have been willing to wait, we consider these trials right-censored. To ensure that the survival curves covered comparable time ranges in both temporal environments, we restricted our analyses to the first 20 seconds of every trial. To account for individual variability, we ran a mixed effects Cox proportional hazards model with random effects for the effect of temporal environment on each participant. Lesion group was modeled as a fixed effect, with the healthy control group as the reference condition. Our predictors of interest were lesion group, temporal environment, and the interaction between the two. Significant positive coefficients for the effect of temporal environment suggest that participants were more likely to skip a trial – that is, to wait less – in the limited persistence condition. Significant positive coefficients for the effect of lesion group suggested that the relevant group waited less than the healthy controls. Finally, significant negative effects for the interaction between lesion group and environment indicate a reduced sensitivity to environmental characteristics compared to healthy controls. From this survival analysis, we also estimated the area under the curve (AUC) using the “desctools” package in R. The AUC is commonly used as proxy for mean survival time and provided us with an estimate of how many of the first 20 seconds participants were, on average, willing to wait.

Finally, we followed the approach proposed in McGuire et al.^14^ to obtain a running estimate of each participants’ quitting threshold across the entire task. The running estimate was initialized to the longest time the participant waited on any trial up to (and including) the first time they quit. Willingness to wait was then updated trial by trial: if a participant waited for longer than the current running estimate, we raised the value to the observed waiting time. If instead they quit at a delay shorter than the current running estimate, we dropped the value to the observed quit time. Finally, if they waited until the end of a trial but the reward came earlier than their current willingness to wait, we did not update the estimate. This procedure allows us to compute a “best guess” measure for participants’ tolerance of waiting throughout the task. To assess whether participants learned to wait longer in the high-persistence condition, we computed a difference score for each participant by subtracting their running estimate of willingness to wait in the limited-persistence condition from their willingness to wait estimate in the high-persistence condition. We then averaged this difference across each minute of the task and assessed whether it was significantly greater than zero for each lesion group.

### Computational Model

To model behavior in the willingness-to-wait task, we fit a Q-learning model that has been adapted for this task.^24^ This model successfully explains behavior in identical willingness-to-wait paradigms, outperforms other competing modeling frameworks like R-learning^26^ and a Q-learning variant with only one learning rate, and yields reliable parameter estimates.

#### Task Representation

We represent the willingness-to-wait task as a Markov decision process (MDP) in which the waiting period is broken into discrete time steps, each 1 second long. Every trial begins at time-step *t* = -2, and a token appears after a 2-second intertrial interval at *t* = 0. For each time step after that, the agent must decide whether to keep waiting or to quit. If the agent chooses to wait and the token remains unmatured, the decision process proceeds to the next time step. Alternatively, if the agent quits or the token matures, the agent returns to *t* = −2 to start a new trial. We will refer to the trial-wise payoff as *R*, with *R* = 0 if the agent does not wait until the token matures, or *R* = 10 if it does. The total number of time steps within a trial is referred to as *T*.

#### Model Description

Broadly, the Q-learning model assumes that the agent compares the value of quitting to the value of waiting at each timestep to arrive at a decision. The value of continued waiting depends on how much time has elapsed since the token appeared and is referred to as *q*(wait, t). Conversely, the value of quitting is not time-dependent, since the result of quitting is the same regardless of how long the waiting period has lasted. The agent makes a wait-or-quit decision every second according to the following choice probability:

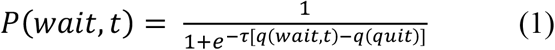

where τ is the inverse temperature parameter that controls action selection noise. Higher values of τ denote more deterministic choices in favor of the higher value option. The closer τ is to zero, the more random an agent’s choices.

At the end of a trial, the value of each preceding time step is updated simultaneously according to a feedback signal *g(t). g(t)* corresponds to a local estimate of the total discounted rewards following time step *t*. Computing it requires adding up the current-trial reward and the prospective reward from future trials, modulated by a discount parameter γ. The current-trial reward corresponds to *R* and is received explicitly at the end of the trial. Conversely, the prospective reward from all future trials is equivalent to the value of quitting, *q(quit)*, and is derived from the agent’s best-guess estimate of prospective rewards given the recurrent nature of the task. Consequently, we have

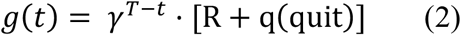

where γ is the discount parameter and *T* − *t* measures the time difference between the end of the trial and the current time step. Trials with positive and non-positive values of *R* are then updated separately as follows:

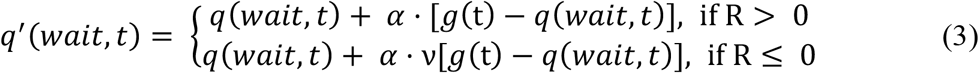

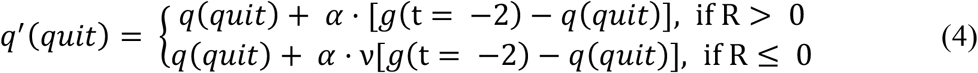

α is a learning rate parameter that determines the step size of each update, while ν is a valence-dependent bias that modulates the degree of updating in cases where a reward was not received. ν < 1 indicates that the agent updates its beliefs more on trials that result in positive feedback than on trials that result in a non-positive payoff (a tendency sometimes referred to as optimism bias^27^). The value of quitting is updated towards the feedback signal of the initial timestep *t* = -2 since quitting leads to beginning again on a new trial.

All that is now missing from the model is how the initial value estimates are determined. *q*(*quit*) is defined as the average optimal rate of return across the HP and LP environments, divided by a scaling constant to reflect discounting of future rewards (we take this discount rate to be equal to 0.85, which roughly matches the empirical median):

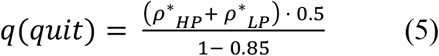

This formulation ensures that the value of quitting is agnostic as to which environment the agent is initially placed in, but nonetheless corresponds to a reasonable range. Finally, the model initializes the value of waiting at individual time steps based on the assumption that there is some upper limit to how much persistence is advantageous. As such, q(*wait, t*) is given linearly decreasing initial values as a function of *t*. A free parameter *η* is added as an intercept term that determines the initial indifference point (i.e., the value of *t* at which q(*wait, t*) drops below the value of quitting):

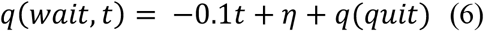

A larger estimate of *η* indicates a greater initial propensity to wait.

#### Model Fitting

We fit the model in Stan^28^ separately for each participant. This yielded posterior distributions of parameter estimates sampled using Markov Chain Monte Carlo (MCMC) with Uniform priors (*τ* ∈ [0.1, 22], α ∈ [0, 0.3], *γ* ∈ [0.7, 1], *η* ∈ [0, 6.5]). We then calculated the mean of the posterior distributions to use as point estimates of parameter values in the subsequent sections.

In order to maximize the reliability of the parameter estimates, we included all the data collected from each participant, including the second block from each session that is excluded from the behavioral analyses. We assumed that participants learned continuously throughout their participation in the experiment, so we did not reinitialize value estimates across sessions or blocks We fit the model with four chains of 8000 samples for each participant, discarding the first 4000 samples as burn-in. If the model did not converge for a given participant (R-hat > 1.01, effective sample size per chain < 100, or divergent transitions detected), we fit the model again with four chains and 20000 samples per chain (10000 as burn-in). If the model still did not converge, we excluded the participant from our modeling analyses. This resulted in excluding 3 participants from the healthy control group, 3 participants from the dmPFC/AI group, 2 participants from the vmPFC group, and 2 participants from the frontal control group.

## Results

### Behavior

If participants are sensitive to the temporal statistics of each condition, they should wait longer in the high persistence (HP) than in the limited persistence (LP) environment. Our experimental design presented the timing environments in a fixed order to prioritize sensitivity to between-group differences; nevertheless, we examined whether the healthy control group data contained within-participant patterns consistent with the previously observed effect of timing environment. To quantify the effect, we compared the mean area under the survival curve (AUC) from the proportional hazards model in each of the two environments. Doing so amounted to assessing how many of the first 20 seconds of each trial participants waited on average, thus factoring out the longer maximum waiting time in the LP condition (40 s in LP versus 20 s in HP). Consistent with previous results^14,15,17–19,29^, we found that the average AUC was significantly greater in the HP than the LP environment in the healthy control group (t(17) = 4.6; p = 0.00024, Figure 2B).

**Figure 2.**
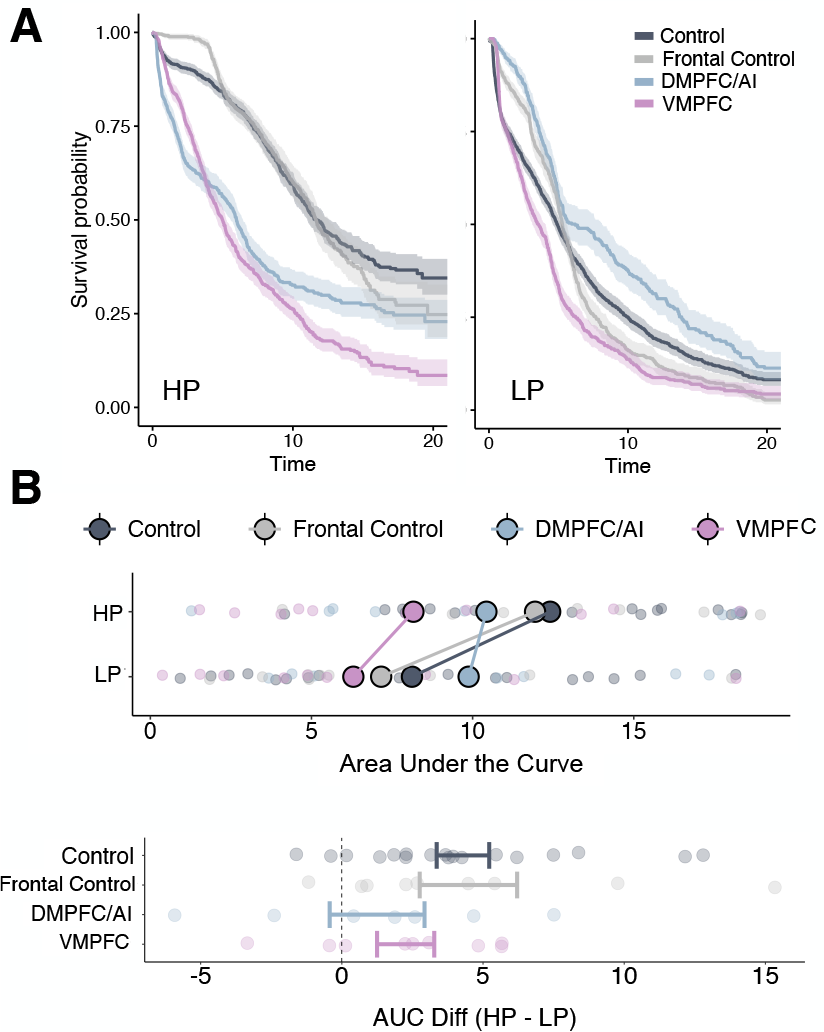
Within-trial willingness to wait for each group of participants with lesions and healthy controls. **(A)** Each group’s mean survival curve is plotted for the high-persistence environment (left) and the limited-persistence environment (right). **(B) Top:** Area under the survival curve (AUC) for each group of participants in the high- and limited-persistence environments. Opaque circles indicate group means and each transparent circle represents a participant. **Bottom:** Difference in average time waited in the HP versus the LP condition, separated by lesion group. Range corresponds to the mean ± the standard error for each group.

The Frontal Control (FC) group exhibited waiting behavior that was quantitatively similar to that of the healthy controls (Figure 2A and B). The average AUC for frontal controls was significantly greater in the HP condition than in the LP condition (difference in AUC: t(8) = 2.5; p = 0.031, Figure 2B). To quantify this similarity, we fit a survival model that predicted time waited as a function of both lesion group and the interaction between lesion group and temporal environment. Neither of the two coefficients showed a significant effect of the FC group relative to healthy controls (β_FC_ = 0.018; p = 0.80; β_FC*env_ = 0.30; p = 0.58). Together, these findings suggest that the waiting behavior exhibited by the frontal controls follows similar principles of dynamic valuation.

The remaining participants showed deficits in their ability to adaptively calibrate wait duration, and their behavior differed significantly from that of the healthy controls (Figure 2A). Although both the vmPFC and dmPFC/AI lesion groups behaved suboptimally, however, they exhibited different biases with respect to their sensitivity to environmental differences versus overall reward. On one hand, participants with lesions to the vmPFC were less willing to wait for the reward overall, and especially when the temporal statistics of the environment favored persistence. The mean AUC for participants with vmPFC lesions was 8.1 ± 5.6 seconds in the HP condition, compared to 12.4 ± 4.4 seconds for the healthy controls (t(26) = 2.1, p = 0.037). Relatedly, the survival term from the proportional hazards model was significantly different for the vmPFC group compared to healthy controls (β_vmPFC_=1.09, p < 0.0001, Figure 2A), suggesting a lower survival rate overall.

Conversely, participants in the dmPFC/AI group did not wait significantly less time than the healthy controls: their average AUC was 10.1 ± 6.3 seconds across the two conditions, which was statistically indistinguishable from the healthy control group’s overall mean of 10.3 ± 5.6 seconds (t(23) = 0.09, p = 0.92). However, participants with lesions to the dmPFC/AI did not calibrate their waiting behavior based on the statistics of the temporal environment. This reduced sensitivity to the difference between the high and limited persistence conditions was evident in the effect on wait duration of the interaction between environment and dmPFC/AI lesion group compared to controls (β_dmPFC/AI*env_ = -1.15; p = 0.049). Similarly, we found no significant difference in AUC across environments for participants in the dmPFC/AI group (t(6) = 0.74, p = 0.48, Figure 2B).

Thus far, we have focused on trial-wise analyses of waiting behavior that collapse across the entire block. While it is clear that lesion location modulates willingness to wait overall, the process by which lesions might affect how temporal environments are learned remains unclear. Figure 3 shows a local estimate of giving-up-time, averaged across 1s timepoints and across participants, as it evolved throughout learning. This estimate is dynamically updated based on participants’ trial-by-trial behavior. By the end of the 12 minutes, both the healthy participants and the frontal controls had learned to wait longer in the high-than the limited-persistence environment, as indexed by a reliably positive difference in willingness to wait in HP versus LP (one-sample t-test averaged over the second half of the task: p = 0.002 and p < 0.001, respectively). The evolution of willingness to wait for the dmPFC/AI and vmPFC groups followed a more unusual pattern, however. For the participants with dmPFC/AI lesions, learning appeared to be somewhat erratic, with no sensitivity to temporal environment by the end of the task (one-sample t-test averaged over the second half of learning: p = 0.84). Though the vmPFC lesions group did, on average, wait significantly longer in the HP than the LP condition in the second half of the task (one-sample t-test: p = 0.02), it is not clear that this difference was the result of learning. Indeed, while the magnitude of the difference in willingness to wait across environments increased with time for both the healthy participants and frontal controls (β = 0.31, p = 0.001 and β = 0.21, p = 0.05, respectively), this progression was not evident for either the dmPFC/AI or vmPFC lesion group (β = 0.28, p = 0.09 and β = -0.02, p = 0.72).

**Figure 3.**
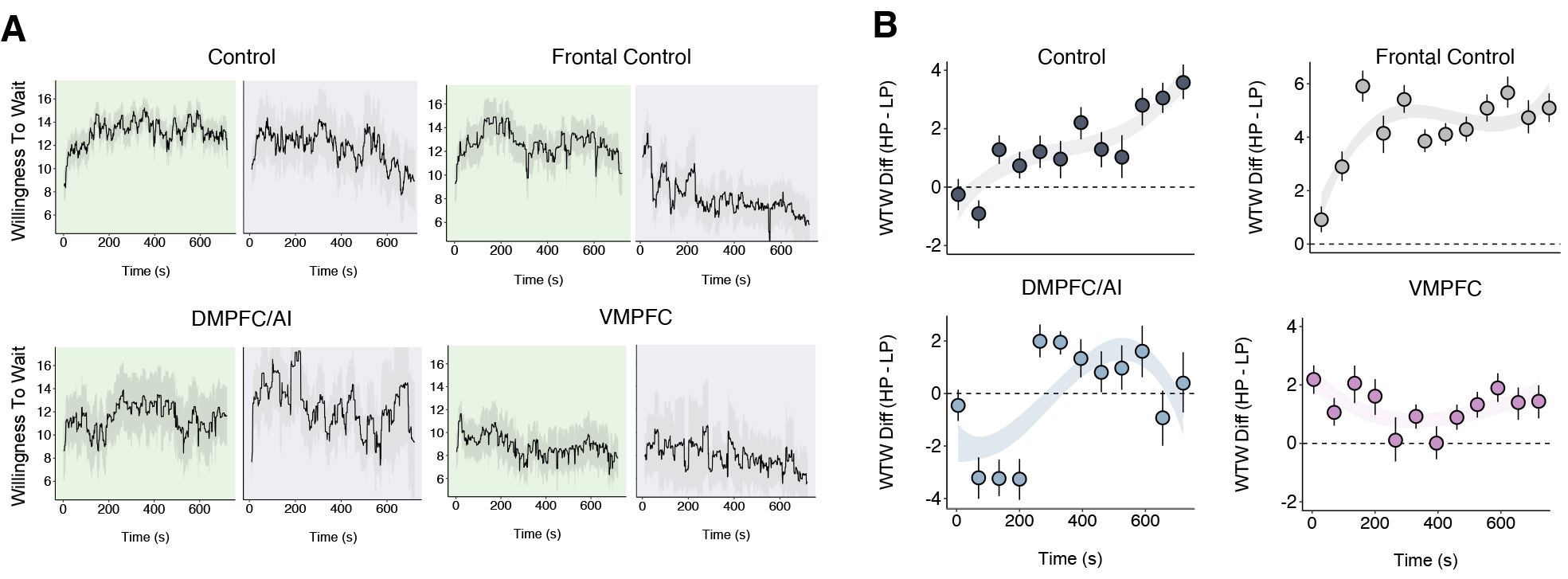
**(A)** Running average (± standard error) of willingness to wait across both blocks of the experiment, sampled at 1 second intervals. **(B)** Evolution of the difference in estimated willingness to wait in the high-persistence vs. limited-persistence condition over time. Smoothed line estimated from a cubic fit to the data, including standard error. Filled circles correspond to mean WTW difference across 72-second bins (10 total bins); lines correspond to bootstrapped confidence intervals around each mean.

### Parameter Comparisons

The model-free analyses we have just described emphasized the functional specificity of different brain regions, with the vmPFC having a role in overall persistence and the dmPFC/AI involved in adaptive calibration. To further understand how different brain regions contribute to waiting behavior, we fit a computational model of willingness to wait to the data, as described in the Methods section and Figure 4A. The model contains five free parameters, which we estimated separately for each individual. Each parameter captures a distinct feature of the decision process and can provide a more mechanistic glimpse into the origin of the behavioral deficits we observe. We ensured that the model could recapitulate behavior by simulating the AUC for each participant based on model-derived estimates of their parameter values, as plotted in Figure 4B. Across both temporal environments, there were no significant group differences between the true behavioral AUC and the AUC derived from model simulations (see Figure 4B for statistical tests). This shows that the model could successfully capture behavioral tendencies as they varied across participants and lesion groups.

**Figure 4.**
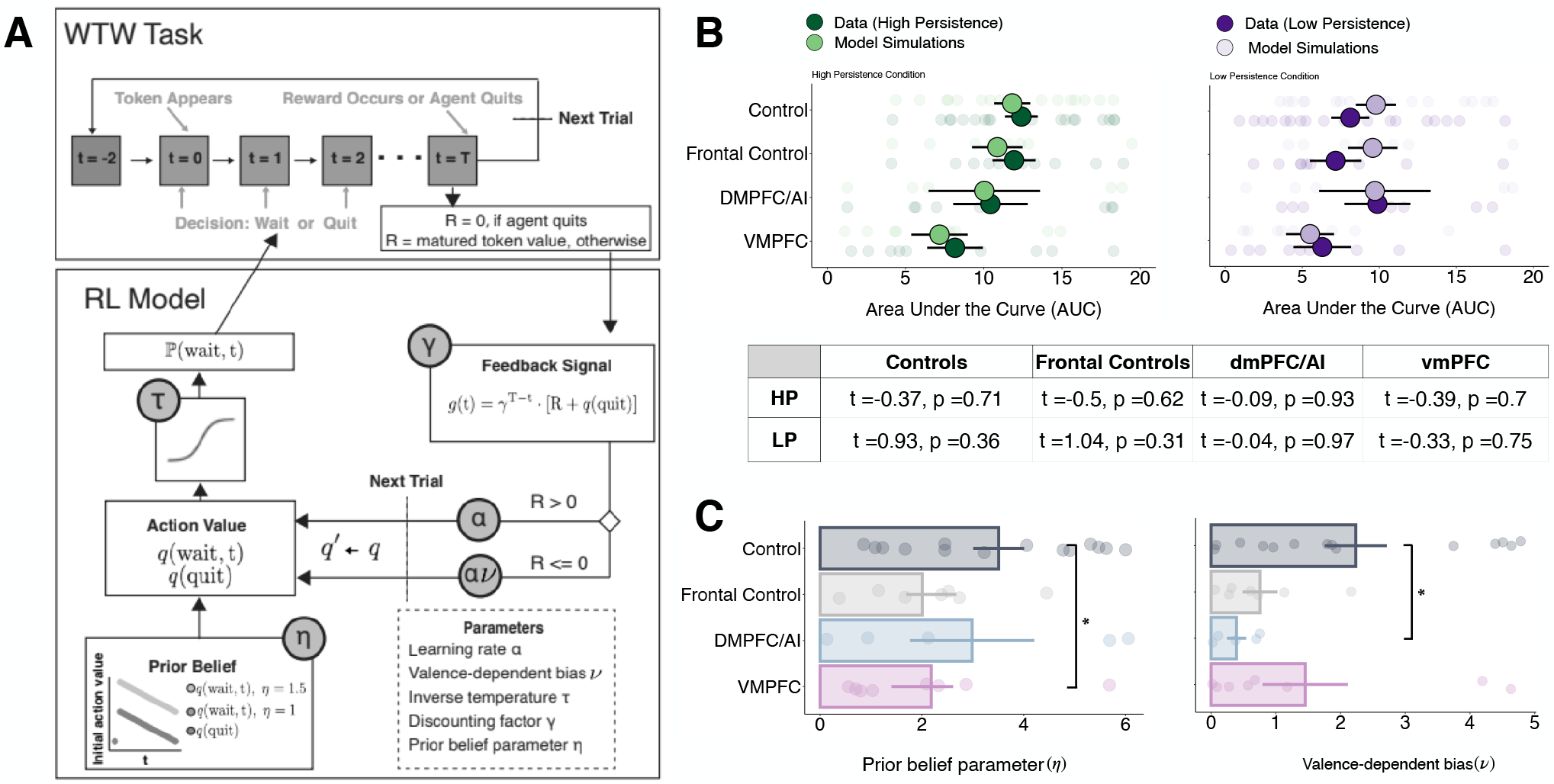
**(A)** Schematic of the 5-parameter reinforcement learning model used to computationally characterize willingness to wait. It assumes that participants maintain estimates of the value of quitting and the value of waiting at each time step and compare the two in order to determine choice. **(B)** Comparison of AUCs from the behavioral data (plotted in darker colors) and model simulations based on the parameter estimates for each participant (plotted in lighter colors). The model can recapitulate the differences in waiting behavior across lesion groups. **(C)** Parameter estimates derived from the computational model for prior belief (left) and valence-dependent bias (right). As a group, participants with vmPFC lesions showed reduced initial willingness to wait, while dmPFC/AI damage led to lower valence-dependent updating.

We found that the model captured the behavioral dissociations outlined above through significant differences in the values of two parameters (Figure 4C). First, people with lesions to the vmPFC had lower values of initial willingness to wait on average, as captured by parameter *η* (Wilcoxon rank sum test: p = 0.047). As described in Equation 6, *η* is an intercept term that determines when the value of waiting falls below the value of quitting for each person, before any learning. Higher values of *η* indicate more patient behavior at baseline, whereas lower values of *η* indicate an initial reluctance to wait. Second, lesions to the dmPFC or anterior insula resulted in lower estimates of the valence-dependent updating parameter ν (Wilcoxon rank sum test: p = 0.014). This difference suggests that participants with dmPFC/AI lesions were less able to learn from quit trials. In fact, the within-group average value of ν was 0.39 (compared to 2.23 for the healthy controls), indicating a strong bias towards greater belief-updating from positive, as opposed to negative or neutral, outcomes. Finally, though we see a marginal difference between the value of ν for healthy versus frontal controls (Wilcoxon rank sum test: p = 0.079), none of the parameters estimated from the model were reliably different between the healthy control and the frontal control groups.

## Discussion

Here, we provide causal evidence for the necessity of the vmPFC and dmPFC/anterior insula in the adaptive calibration of persistence. While participants with lesions to the vmPFC were less willing to wait for delayed rewards overall, lesions to the dmPFC and anterior insula caused a deficit in people’s ability to learn from quit trials. Crucially, lesions to lateral regions of the prefrontal cortex, an area of the brain implicated in self-control^5,30–32^, did not affect performance. We validated these findings with a computational model of task performance, which highlighted deviations in initial willingness to wait for the vmPFC group and decreases in the rate of learning from negative feedback for the dmPFC/AI group. Thus, both neural substrates contribute necessary and functionally distinct computations to adaptive persistence.

Our finding that lesions to the vmPFC led to reduced willingness to wait across both temporal environments echoes its known role in the representation of subjective value.^20–22^ The subjective nature of this representation is highlighted by the fact that vmPFC activity during the willingness-to-wait task is modulated by temporal context and dynamically changes over the course of the delay interval.^15^ However, we did not find that lesions to the vmPFC specifically disrupted the calibration of waiting behavior to the temporal environment. Rather, we found that lesions caused participants to wait less overall. This result suggests that the vmPFC may play a more general role in maintaining behavior towards delayed rewards, which is in line with previous findings in intertemporal choice tasks. Participants with mPFC lesions exhibit a stronger preference for smaller-sooner reward options, and cortical thickness in the vmPFC and neighboring orbitofrontal cortex (OFC) is correlated with patience in a delayed discounting paradigm.^33,34^ Thus, the vmPFC may play an important role in tempering impulsivity by holding a neural representation of the upcoming reward. This framework is also in line with reports of increased impulsivity in people that sustain damage to the vmPFC.^35,36^ VmPFC damage appears to bias people away from considering the long-term consequences of behavior, simultaneously leading to more present-oriented behavior^37^ and – in our experimental context – decreased willingness to wait.

Our finding that lesions to the anterior insula and dmPFC lead to impairments in learning from feedback is consistent with existing evidence that these brain regions play a role in belief updating. For instance, McGuire et al.^38^ found reliable activity in the insula and dmPFC in a change-point detection task that required participants to update their beliefs about reward contingencies. That work identified increased activity in these brain regions when a surprising outcome indicated that participants needed to revise their mental model of the task. Similarly, Stöttinger et al.^39^ showed that participants with insular lesions struggle on tasks that required the updating of mental models across perceptual and non-perceptual domains. These findings suggest that the insula and the dmPFC play a necessary role in the selective integration of task-relevant information, which could be related to their involvement in the brain’s “salience network”.^40^

In the context of the willingness to wait task, McGuire and Kable^15^ highlighted a role for the insula and dmPFC in signaling imminent quit decisions. If we conceptualize the last couple of seconds before a quit decision as being particularly salient, then this is in line with the network’s role in flagging relevant information. Furthermore, it may be that the signal in the dmPFC and anterior insula that ramps up right before a quit decision helps the brain to learn from quit events in addition to rewards. Although this interpretation is speculative, it is supported by the results from the computational model; the dmPFC/AI lesion group differed specifically in a parameter that governed learning from quit trials.

So far, we have emphasized the role vmPFC and dmPFC/anterior insula play in the calibration of persistence. Of equal importance, however, is the finding that participants in the frontal control group – who have lesions to more lateral portions of the prefrontal cortex – were not impaired in the willingness to wait task. Self-control is often thought to rely on lateral regions of the prefrontal cortex, and especially the inferior frontal gyrus (IFG). For instance, Heatherton et al. (2011) put forth a framework in which top-down control from the lateral PFC and the IFG over subcortical areas like the striatum and the amygdala is necessary for successful self-regulation. This theory is inspired by evidence that response inhibition, or the ability to override a prepotent response in favor of the correct choice, relies on lateral PFC activation.^5,41–43^ In our results, however, we see no difference in waiting behavior between participants with lesions to the lateral regions of the prefrontal cortex and healthy controls. If choosing to wait were really a matter of sustained self-discipline, we would expect lesions to the lateral PFC to result in decreased waiting overall, rather than the intact performance we observe.

To support our claims about functionally distinct roles of different regions of the frontal cortex in voluntary persistence, we fit a task-specific reinforcement learning model to the data. This model has previously been validated on datasets with much larger sample sizes than ours,^24^ following guidelines for the computational modeling of behavioral data outlined in Wilson and Collins (2019).^44^ We are thus confident that the model can capture meaningful differences across lesion groups, particularly in light of the clear behavioral deficits we observe. Nonetheless, in line with the general notion that all models are at least partially wrong, we recognize that many other well-suited modeling alternatives exist. Of particular interest here is the distinction between model-free and model-based reinforcement learning. While the Q-learning model we implemented here is model-free in that it does not assume that people are learning higher-level distributions of externally imposed delay durations, a model-based alternative might assume that participants are forming beliefs about the temporal statistics of each environment^45^. Although this is a reasonable assumption, we recently showed that providing participants with explicit descriptions of the wait durations did not significantly improve their performance^18^. Thus, learning in this paradigm may not rely on model-based processing. Formalizing a computational alternative to model-free Q-learning will help us further examine this issue in the future.

With this study, we have provided evidence that distinct regions of the brain play different causal roles in determining how long people are willing to wait for future rewards. Although we focus here on participants with brain lesions, we believe that the willingness to wait task and the broader framework of persistence as valuation is relevant to other clinical domains. Indeed, mental illnesses like anxiety and depression often involve deviations in how reward is processed, particularly when its receipt is delayed or uncertain^46–48^. As such, we expect that clinical populations may exhibit systematically different choice behavior in the paradigm we have described. For instance, Mukherjee et al. (2020) administered the willingness to wait task in a sample of participants with major depression and found that they waited significantly less long in the high persistence environment than healthy controls. In the future, we hope to continue this line of work in other clinical populations, leveraging what we know about the computations that are required for adaptive behavior to shed light on processes that may go awry in mental illness.

## Acknowledgments

We would like to thank Lesley Fellows for facilitating access to participants in Montreal, Anjan Chatterjee and Roy Hamilton for facilitating access to participants in Philadelphia, Christine Déry and Eileen Cardillo for coordinating participants in Montreal and in Philadelphia, and all of the participants themselves, without whom this work would not have been possible. This work was supported by National Institute on Drug Abuse (NIDA) R01-DA029149 to JWK, National Institutes of Health (NIH) F32-DA030870, National Science Foundation (NSF) BCS-1755757, and National Institutes of Health (NIH) R21-MH124095 to JTM. This research was supported in part by the Intramural Research Program of the NIH, NIDA (ZIA DA000642). The content of this paper is solely the responsibility of the authors and does not reflect the official views of the NIH.

